# Pain-related fear – Dissociable neural sources of different fear constructs

**DOI:** 10.1101/251751

**Authors:** Michael Lukas Meier, Andrea Vrana, Barry Kim Humphreys, Erich Seifritz, Philipp Stämpfli, Petra Schweinhardt

## Abstract

Fear of pain demonstrates significant prognostic value regarding the development of persistent musculoskeletal pain and disability. Its assessment often relies on self-report measures of pain-related fear by a variety of questionnaires. However, based either on “fear of movement/(re)injury/kinesiophobia”, “fear avoidance beliefs” or “pain anxiety”, pain-related fear constructs seemingly differ while the potential overlap of the questionnaires is unclear. Furthermore, the relationship to other anxiety measures such as state or trait anxiety remains ambiguous. Because the neural bases of fearful and anxious states are well described, advances in neuroimaging such as machine learning on brain activity patterns recorded by functional magnetic resonance imaging might help to dissect commonalities or differences across pain-related fear constructs. We applied a pattern regression approach in 20 non-specific chronic low back pain patients to reveal predictive relationships between fear-related neural information and different pain-related fear questionnaires. More specifically, the applied Multiple Kernel Learning approach allowed generating models to predict the questionnaire scores based on a hierarchical ranking of fear-related neural patterns induced by viewing videos of potentially harmful activities for the back. We sought to find evidence for or against overlapping pain-related fear constructs by comparing the questionnaire prediction models according to their predictive abilities and associated neural contributors. The results underpin the diversity of pain-related fear constructs by demonstrating evidence of non-overlapping neural predictors within fear processing regions. This neuroscientific approach might ultimately help to further understand and dissect psychological pain-related fear constructs.

**Significance:** Pain-related fear, often assessed through self-reports such as questionnaires, has shown prognostic value and clinical utility for a variety of musculoskeletal pain disorders. However, it remains difficult to determine a common underlying construct of pain-related fear due to several proposed constructs among questionnaires. The current study describes a novel neuroscientific approach using machine learning of neural patterns within the fear circuit of chronic low back pain patients that has the potential to identify neural commonalities or differences among the various pain-related fear constructs. Ultimately, this approach might afford a deeper understanding of the suggested constructs and might be also applied to other domains where ambiguity exists between different psychological constructs.

## 1. Introduction

Self-report measures of emotional states are paramount for behavioral neuroscience by enabling the understanding of brain response patterns (Shrout et al., 2017). However, the validity of self-reports is limited (Choi and Pak, 2005), probably also because often overlapping psychological constructs are assessed, illustrated by the fact that various questionnaires attempt to assess related constructs. One example is pain-related fear (PRF), which is a major explanatory variable of disability in patients with persistent musculoskeletal pain (Crombez et al., 1999; Vlaeyen and Linton, 2000; Vlaeyen et al., 2016). Back straining activities (i.e. bending and lifting) are the most feared and pain-provoking movements among people with low back pain (LBP), based on ratings of perceived harmfulness or physiological responses (Caneiro et al., 2017; Leeuw et al., 2007a; Stevens et al., 2016; Glombiewski et al., 2015). As such, active or passive (e.g. through pictures) bending and lifting have been frequently used to provoke PRF (Leeuw et al., 2007c; Caneiro et al., 2017; Barke et al., 2016; Trost et al., 2009). For the assessment of PRF, various questionnaires exist based on constructs such as fear of movement/(re)injury/kinesiophobia, fear avoidance beliefs or pain anxiety. However, despite the clinical relevance of PRF self-reports, their construct validity remains ambiguous and there is an open debate on what their scores reflect on the fear-anxiety spectrum (Lundberg et al., 2011; Caneiro et al., 2017). Fear represents a reaction to an imminent threat, preparing the individual for “fight-flight-freeze”, whereas anxiety is described as more diffuse (e.g. cognitions about a future threat) (Kreddig and Hasenbring, 2017; LeDoux and Pine, 2016). While PRF questionnaires do not clearly distinguish between these emotions (Kreddig and Hasenbring, 2017; Lundberg et al., 2011), brain research provides evidence for a functional differentiation of fear and anxiety. Both emotions are controlled by the fear circuit (Tovote et al., 2015), however, subcortical regions (e.g. the amygdala) seem to be more involved in fast and defensive fear reactions (short defensive distance to threat) while cortical regions (e.g. the prefrontal cortex) are more likely responsible for complex cognitions of fear and anxiety (large defensive distance to threat) (Qi et al., 2018; McNaughton and Corr, 2004). Therefore, advances in neuroimaging enable exploring the (sub-)cortical contributions to PRF constructs through examining interrelations between self-reported emotional states and brain response patterns. Specifically, machine learning techniques such as multivariate pattern analysis (MVPA) if applied to functional magnetic resonance imaging (fMRI) data make it possible to directly study the predictive relationship between a content-selective cognitive or emotional state (expressed as a label) and corresponding multivoxel fMRI activity patterns (Haynes, 2015; Hebart and Baker, 2017). The label may have discrete (classification) or continuous (regression) values such as questionnaire scores (Formisano et al., 2008). Therefore, we provoked PRF by presenting video clips of daily activities including bending and lifting (harmful condition for the back) and harmless activities such as walking (harmless condition) in a sample of 20 non-specific chronic LBP patients. We applied a pattern regression analysis in combination with Multiple Kernel Learning to assess potential neural predictors of the various PRF constructs based on weighting of 1) harmful and harmless conditions (condition weights) and 2) pattern information within sub-cortical and cortical fear processing regions (region weights). We first contrasted the different PRF questionnaires in terms of their model performance, namely the model’s ability to predict the questionnaires scores based on brain response patterns across fear processing regions. Second, we compared the different prediction models according to the distributions of their condition and region weights to explore potential neural commonalities or differences of related PRF constructs. If the PRF questionnaires share overlapping PRF constructs, then the regions weights should be similarly distributed across fear processing regions. Conversely, if the contributing brain regions vary across the predictions models, this would provide evidence against a common PRF construct across questionnaires. Ultimately, this approach might help to further understand and dissect the various PRF constructs in chronic LBP.

## 2. Methods

### 2.1 Patients

The study was approved by the Ethics Committee Zurich (Switzerland) and all patients provided written informed consent before participation. The study was conducted in accordance with the Declaration of Helsinki and involved a total of 20 patients (mean age = 39.35 years, SD = 13.97 years, 7 females) with non-specific chronic LBP (Table 1). Non-specific chronic LBP is the most common form of back pain (about 85% of the cases) and constitutes a heterogeneous and complex biopsychosocial condition without a specific nociceptive cause (Maher et al., 2017; Deyo and Weinstein, 2001). Patients were recruited via local chiropractic and physiotherapy centres as well as via online advertisements. Inclusion criteria were low back pain of at least 6 months duration and age between 18 and 65 years. Exclusion criteria were a history of psychiatric or neurological disorders and specific causes for the pain (e.g. infection, tumour, fracture, inflammatory disease) that were ruled out by an experienced clinician.

**Table 1.** Patient characteristics and descriptive statistics of questionnaires. Tampa Scale of Kinesiophobia (TSK, SF = somatic focus subscale, AA = activity avoidance subscale), Pain Anxiety Symptome Scale (PASS, PASSc = cognitive anxiety, PASSe = escape/avoidance, PASSf = fear, PASSp = physiology), Fear Avoidance Beliefs (FABQ, FABQ-PA = physical activity, FABQ-W = work), State-Trait Anxiety Inventory (S-Anxiety, T-Anxiety).

|  | *Minimum* | *Maximum* | *Mean* | *SD* |
| --- | --- | --- | --- | --- |
| ***cLBP patients (N = 20, 7 females)*** |  |  |  |  |
| *Age* | 21 | 62 | 39,35 | 13,97 |
| *TSK-17* | 26 | 52 | 36,90 | 5,59 |
| *TSK-13* | 16 | 43 | 27,60 | 5,96 |
| *TSK-11* | 13 | 38 | 23,20 | 5,71 |
| *TSK-11-SF* | 5 | 16 | 9,70 | 2,69 |
| *TSK-11-AA* | 5 | 20 | 11,90 | 3,35 |
| *PASS* | 13 | 68 | 38,15 | 16,57 |
| *PASS-C* | 1 | 15 | 8,70 | 4,19 |
| *PASS-E* | 3 | 21 | 9,85 | 4,77 |
| *PASS-F* | 2 | 20 | 9,45 | 5,28 |
| *PASS-P* | 0 | 15 | 7,35 | 4,21 |
| *FABQ* | 3 | 83 | 35,45 | 22,53 |
| *FABQ-PA* | 2 | 21 | 12,80 | 5,59 |
| *FABQ-W* | 0 | 40 | 15,50 | 12,12 |
| *S-Anxiety* | 36 | 53 | 43,70 | 4,78 |
| *T-Anxiety* | 31 | 59 | 43,00 | 6,05 |
| *PainDETECT current pain* | 0 | 8 | 3,77 | 2,49 |
| *PainDETECT strongest pain* | 2 | 10 | 6,15 | 2,16 |
| *PainDETECT average pain*<br>*(previous 4 weeks)* | 1 | 7 | 3,75 | 1,88 |
| *Ratings harmful activities* | 0 | 10 | 5,44 | 2,38 |
| *Ratings harmless activities* | 0 | 5 | 1,28 | 1,32 |

### 2.2 Self-report measures of pain-related fear

PRF was assessed using several questionnaires:

(1) The Tampa Scale of Kinesiophobia questionnaire (TSK) (Vlaeyen et al., 1995; Kori et al., 1990) was used to assess fear of movement/(re)injury and kinesiophobia. The 17-item German version of the TSK (TSK-17) with satisfactory internal consistency (Cronbach’s α = 0.76–0.84) contains statements focusing on fear of physical activity rated on a 4-point Likert scale from 1 = “strongly disagree” to 4 = “strongly agree” (Rusu et al., 2014). Due to additional versions of original 17-item TSK questionnaire, we also calculated the questionnaire scores of the 13- and 11-item TSK versions (TSK-13, TSK-11). The 13-and 11-item versions were previously validated by confirmatory factor analysis and demonstrated acceptable levels of internal consistency (Cronbach’s α =0.80) (Goubert et al., 2004; Tkachuk and Harris, 2012). A two-factor solution of the TSK-11 version provides the best fit in terms of explaining variance across German, Dutch, Swedish and Canadian samples and included the subscales “activity-avoidance” (TSK-AA, the belief that that activity may result in (re)injury or stronger pain) and “somatic focus” (TSK-SF, the belief in underlying and serious medical problems) (Roelofs et al., 2007; Rusu et al., 2014).

(2) The German version of the fear avoidance beliefs questionnaire (FABQ) (Waddell et al., 1993; Pfingsten et al., 2000) consists of 16 back pain-specific items related to fear avoidance beliefs rated on a 7-point rating scale (0 = “completely disagree” to 6 = “completely agree”). It includes two distinct and established subscales related to beliefs about on how work (FABQ-W) and physical activity (FABQ-PA) affects LBP with internal consistencies of α = 0.88 and α = 0.77, respectively (Waddell et al., 1993).

(3) The short version of the Pain Anxiety Symptoms Scale (PASS-20) assesses fear and anxiety responses related to pain including cognitive, physiological and motor response domains (McCracken and Dhingra, 2002). Items on the PASS-20 are measured on a 6-point Likert scale and relate to four different subscales including cognitive anxiety (PASS-C), fear (PASS-F), physiology (PASS-P) and escape/avoidance (PASS-E) (Roelofs et al., 2004b). The German version of the PASS-20 has an internal consistency of α = 0.90 (Kreddig et al., 2015).

Furthermore, patients were asked to fill out the painDETECT (PD-Q) questionnaire that includes three 11-point numeric rating scales (NRS), with 0 being “no pain” and 10 being the “worst imaginable pain” to assess current pain, strongest and average pain intensity in the previous 4 weeks (Freynhagen et al., 2006). Finally, to investigate potential differences or shared variance between PRF and general anxiety, we used the State-Trait Anxiety Inventory (STAI), the most widely used self-report measure of anxiety which including two subscales (Spielberger and Gorsuch, 1983; Julian, 2011): The State Anxiety Scale (S-Anxiety) assesses the current state of anxiety whereas the Trait Anxiety Scale (T-Anxiety) evaluates more stable aspects of anxiety such as “anxiety proneness” (Julian, 2011). All questionnaires were administered at the fMRI appointment prior to brain scanning. We tested the scores of the different questionnaires for the assumption of normality of the data using the Shapiro-Wilk test and visually using Q-Q plots implemented in IBM SPSS Statistics (version 23) (Ghasemi and Zahediasl, 2012).

### 2.3 Scanning protocol and design

Brain imaging was performed on a 3-T whole-body MRI system (Philips Achieva, Best, Netherlands), equipped with a 32-element receiving head coil and MultiTransmit parallel RF transmission. Each imaging session started with a survey scan, a B1 calibration scan (for MultiTransmit), and a SENSE reference scan. High resolution anatomical data were obtained with a 3D T1-weighted turbo field echo scan consisting of 145 slices in sagittal orientation with the following parameters: FOV = 230 × 226 mm^2^; slice thickness = 1.2 mm; acquisition matrix = 208 × 203 (resulting in a voxel resolution of 1.1mm x 1.1mm x 1.2mm); TR = 6.8 ms; TE = 3.1 ms; flip angle = 9°; number of signal averages = 1. Functional time series were acquired using whole-brain gradient-echo echo planar imaging (EPI) sequences (365 volumes), consisting of 37 slices in the axial direction (AC-PC angulation) with the following parameters: field of view (FOV) = 240 × 240 mm^2^; acquisition matrix = 96 × 96; slice thickness = 2.8 mm (resulting in a voxel resolution of 2.5mm x 2.5mm x 2.8mm); interleaved slice acquisition; no slice gap; repetition time (TR) = 2100 ms; echo time (TE) = 30 ms; SENSE factor = 2.5; flip angle 80°.

The PRF-provoking stimuli (harmful condition) consisted of video clips with a duration of 4 s recorded from a 3rd person perspective (Meier et al., 2016). The video clips showed potentially harmful activities (back straining movements such as bending and lifting) selected from the Photograph Series of Daily Activities (PHODA) (Leeuw et al., 2007a). The original PHODA was developed in close collaboration with human movement scientists, physical therapists, and psychologists and is comprised of a fear hierarchy based on ratings of perceived harmfulness of daily activities in patients with chronic LBP. From the 40 potentially harmful activities included in the short electronic PHODA version (Leeuw et al., 2007a), we chose three scenarios from the top six most harmful activities, namely shoveling soil with a bent back, lifting a flowerpot with slightly bent back and vacuum cleaning under a coffee table with a bent back. Furthermore, we created video clips of three activities rated as less harmful, such as walking up and down the stairs and walking on even ground (harmless condition). Presentation® software (Neurobehavioral Systems, Davis, CA, USA) was used to present the video clips in a pseudo-randomized order (no more than two identical consecutive trials). The patients were asked to carefully observe the video clips which were displayed using MR-compatible goggles (Resonance Technology, Northridge, CA, USA). The three harmful and harmless activities were each presented five times (30 trials total). After the observation of each video clip, the patients were asked to rate the perceived harmfulness of the activity on a visual analog scale (VAS) which was anchored with the endpoints “not harmful at all” (0) and “extremely harmful” (10). All ratings were performed using a MR compatible track ball (Current Designs, Philadelphia, PA, USA). After the VAS rating, a black screen with a green fixation cross appeared (duration jittered between 6 and 8s). This experimental protocol has been shown suitable for investigations of neural correlates of PRF self-reports in previous fMRI studies based on mass-univariate analyses (Meier et al., 2016; Meier et al., 2017).

### 2.4 MR data organization and pre-processing

We used an existing fMRI dataset of previously reported studies (Meier et al., 2016; Meier et al., 2017). The fMRI data were organized according to the Brain Imaging Data Structure (BIDS), which provides a consensus on how to organize data obtained in neuroimaging experiments. Preprocessing was performed using FMRIPREP (version 1.0.0-rc2, https://github.com/poldracklab/fmriprep), a Nipype based tool (Gorgolewski et al., 2011), which requires minimal user input and provides easily interpretable and comprehensive error and output reporting. This processing pipeline includes state-of-the-art software packages for each phase of preprocessing (see https://fmriprep.readthedocs.io/en/stable/workflows.html for a detailed description of the different workflows). Each T1-weighted (T1w) volume was skullstripped using antsBrainExtraction.sh v2.1.0 (using OASIS template). The skullstripped T1w volume was co-registered to skullstripped ICBM 152 Nonlinear Asymmetrical MNI template version 2009c using nonlinear transformation implemented in ANTs v2.1.0 (Avants et al., 2008). Functional data were slice time corrected using AFNI (Cox, 1996) and motion corrected using MCFLIRT v5.0.9 (Jenkinson et al., 2002). This was followed by co-registration to the corresponding T1w volume using boundary based registration 9 degrees of freedom - implemented in FreeSurfer v6.0.0 (Greve and Fischl, 2009). Motion correcting transformations, T1w transformation and MNI template warp were applied in a single step using antsApplyTransformations v2.1.0 with Lanczos interpolation. Three tissue classes were extracted from T1w images using FSL FAST v5.0.9 (Zhang et al., 2001). Voxels from cerebrospinal fluid and white matter were used to create a mask used to extract physiological noise regressors using aCompCor (Behzadi et al., 2007). The mask was eroded and limited to subcortical regions to limit overlap with grey matter and six principal components were estimated. Independent component analysis (ICA)-based Automatic Removal Of Motion Artifacts (AROMA) was used to generate aggressive motion-related noise regressors. The AROMA classifier identifies motion components with high accuracy and robustness and is superior to motion artefact detection using 24 motion parameters or spike regression (Pruim et al., 2015). Finally, to preserve high spatial frequency while reducing noise, spatial smoothing with a full width at half maximum 4mm Gaussian kernel was applied. To accelerate data pre-processing we performed parallel computing using the Docker environment (https://www.docker.com/) and the GC3Pie framework (https://github.com/uzh/gc3pie) on the ScienceCloud supercomputing environment at the University of Zurich (S3IT, https://www.s3it.uzh.ch/).

### 2.5 MVPA input data

The pre-processed data were subsequently passed to Statistical Parametric Mapping software package (SPM12, version 6906, http://www.fil.ion.ucl.ac.uk/spm/) for model computation using a general linear model (GLM). For each patient a design matrix was built including the onsets of the video clips with a duration of 4s (harmful / harmless activities, each pooled across the three different activities resulting in 15 harmful and 15 harmless stimuli) as separate regressors. In addition, and for each patient, the following nuisance regressors were implemented in the GLM model: (1) the six regressors derived from the component based physiological noise correction method (aCompCor) and (2) the motion-related regressors generated by AROMA (see section 2.4). A high-pass filter with a cut-off of 128 s was used to remove low-frequency noise. Trials were modeled as boxcar regressors and convolved with the standard canonical hemodynamic response function (HRF) as implemented in SPM12. Finally, for each patient, parameter estimates (beta images) for each condition were computed and served as the input images for the MVPA.

### 2.6 Multivariate pattern analysis (MVPA)

Compared to univariate analyses, MVPA can achieve greater sensitivity and is able to detect subtle and spatially distributed effects (Schrouff et al., 2013; Haynes, 2015). A pattern of activity can represent many more different states than each voxel individually, which leads to an information-based view compared to the activation-based view of univariate analyses (Hebart and Baker, 2017). MVPA was performed using routines implemented in PRoNTo v.2.0 (Schrouff et al., 2013). For the read-out of multivariate neural information that might serve as a potential score estimator of the different PRF questionnaires, we applied a newly introduced pattern regression approach based on supervised machine learning and testing phases using Multiple Kernel Learning (MKL). In brief, the objective in supervised pattern recognition regression analysis is to learn a function from data that can accurately predict the continuous values (labels), i.e. f(xi)=yi from a given dataset D={xi, yi}, i=1…N where xi represents pairs of samples or vectors and yi the different labels. Ultimately, the learned function from the learning set is used to predict the labels from new and unseen data (Schrouff et al., 2013). MKL allows to account for brain anatomy (determined by a brain atlas, see section 2.7) and different modalities (such as anatomical/functional data or in the current approach: conditions) during the model estimation by considering each brain region and modality as separate kernels. This approach allows determining the contribution of each brain region (region weights) and condition (condition weights) to the final decision function of the model in a hierarchical manner by simultaneously learning and combining the different linear kernels that are based on support vector machines (SVM) (Rakotomamonjy et al., 2008; Fernandes et al., 2017; Schrouff et al., 2018). Compared to conventional MVPA methods based on whole-brain voxel weight maps, this procedure provides a straight-forward approach to draw inferences on the region level without the need for multiple comparison correction (Schrouff et al., 2018). To account for possible differential contributions of the harmful and harmless conditions to the decision function, we included the individual SPM beta images of each condition as separate modalities in the MKL model (condition weights). The kernels were mean centered and normalized (to account for the different sizes of the involved brain regions) using standard routines implemented in PRoNTo. Subsequently, for each questionnaire, we trained a separate MKL regression model with the respective labels (FABQ, TSK-17-, -13- and -11-item, PASS and all subscale scores, state and trait anxiety). This resulted in a total of 15 MKL models providing outputs for model evaluation, including model performance, region and conditions weights. Furthermore, we trained MKL regression models based on the harmfulness ratings collected during the fMRI measurements (mean ratings of the harmful condition and harmless condition, respectively). To reduce the risk of overfitting for each model, we applied a nested cross-validation procedure using a “leave-one-subject-out” cross-validation scheme to train the model including optimization of the model’s hyperparameter “C” (range [0.1 1 10 100 1000]). Furthermore, to generate a data-based null distribution of the performance measures (r and nMSE, see section 2.8), 1000 permutations (permuting the labels across patients) were computed for each model. Results were considered significant at a threshold of p < 0.05. Finally, the MKL currently implemented in PRoNTo operates with sparsity (L1 regularization) in kernel combinations and might therefore not select brain regions that are highly associated with each other and the prediction variable (these regions will have kernel weights of zero) (Fernandes et al., 2017; Schrouff et al., 2013). This might influence the selection of regions across the models. Therefore, to confirm a dissociation regarding the selected brain regions across the predictive models, we performed a secondary cross-validation by choosing the regions contributing most to the prediction (>10%, see Table 3) of each significant questionnaire model as a separate predictive brain set and subsequently trained and tested the labels of each questionnaire on the predictive brain set of the other models. In doing so, related non-significant results of model performance would reinforce a dissociation of contributing brain regions between the different models and therefore would be indicative of non-overlapping fear constructs.

**Table 2.** Spearman’s rank correlations between the different pain-related fear questionnaires. Tampa Scale of Kinesiophobia (TSK, SF = somatic focus subscale, AA = activity avoidance subscale), Pain Anxiety Symptom Scale (PASS = total score, PASS-C = cognitive anxiety, PASS-E = escape/avoidance, PASS-F = fear, PASS-P = physiology), Fear Avoidance Beliefs (FABQ = total score, FABQ-PA = physical activity, FABQ-W = work), State and Trait Anxiety Inventory (S-Anxiety, T-Anxiety). **p < 0.005 (bold), *p<0.05 (bold).

|  |  | *TSK-17* | *TSK-13* | *TSK-11* | *TSK-11_SF* | *TSK-11_AA* | *PASS* | *PASS-C* | *PASS-E* | *PASS-F* | *PASS-P* | *FABQ* | *FABQ-PA* | *FABQ-W* | *S-ANXIETY* | *T-ANXIETY* | *Rating harmful* | *Rating harmless* |
| --- | --- | --- | --- | --- | --- | --- | --- | --- | --- | --- | --- | --- | --- | --- | --- | --- | --- | --- |
| *TSK-17* | Corr.coeff. | 1,000 | **,834\*\*** | **,800\*\*** | **,609\*\*** | **,759\*\*** | **,611\*\*** | **,503\*** | **,614\*\*** | **,556\*** | **,647\*\*** | ,337 | **,494\*** | ,280 | **,449\*** | ,299 | ,133 | ,289 |
|  | Sig. |  | **,000** | **,000** | **,004** | **,000** | **,004** | **,024** | **,004** | **,011** | **,002** | ,146 | **,027** | ,231 | **,047** | ,200 | ,577 | ,216 |
| *TSK-13* | Corr.coeff. | **,834\*\*** | 1,000 | **,960\*\*** | **,754\*\*** | **,789\*\*** | **,686\*\*** | **,559\*** | **,777\*\*** | **,666\*\*** | **,558\*** | ,344 | **,442\*** | ,339 | **,451\*** | ,139 | ,240 | ,168 |
|  | Sig. | **,000** |  | **,000** | **,000** | **,000** | **,001** | **,010** | **,000** | **,001** | **,011** | ,138 | **,050** | ,144 | **,046** | ,558 | ,307 | ,479 |
| *TSK-11* | Corr.coeff. | **,800\*\*** | **,960\*\*** | 1,000 | **,793\*\*** | **,779\*\*** | **,685\*\*** | **,559\*** | **,766\*\*** | **,643\*\*** | **,565\*\*** | ,360 | ,404 | ,378 | **,470\*** | ,071 | ,276 | ,091 |
|  | Sig. | **,000** | **,000** |  | **,000** | **,000** | **,001** | **,010** | **,000** | **,002** | **,009** | ,120 | ,077 | ,101 | **,037** | ,766 | ,238 | ,703 |
| *TSK-11_SF* | Corr.coeff. | **,609\*\*** | **,754\*\*** | **,793\*\*** | 1,000 | ,350 | **,519\*** | **,462\*** | **,529\*** | **,502\*** | **,502\*** | ,351 | ,315 | ,411 | **,629\*\*** | ,034 | ,044 | ,178 |
|  | Sig. | **,004** | **,000** | **,000** |  | ,131 | **,019** | **,040** | **,016** | **,024** | **,024** | ,130 | ,176 | ,071 | **,003** | ,886 | ,854 | ,452 |
| *TSK-11_AA* | Corr.coeff. | **,759\*\*** | **,789\*\*** | **,779\*\*** | ,350 | 1,000 | **,477\*** | ,268 | **,564\*\*** | **,486\*** | ,421 | ,336 | ,375 | ,280 | ,284 | ,035 | ,270 | ,135 |
|  | Sig. | **,000** | **,000** | **,000** | ,131 |  | **,034** | ,254 | **,010** | **,030** | ,065 | ,147 | ,103 | ,231 | ,224 | ,883 | ,250 | ,570 |
| *PASS* | Corr.coeff. | **,611\*\*** | **,686\*\*** | **,685\*\*** | **,519\*** | **,477\*** | 1,000 | **,895\*\*** | **,899\*\*** | **,886\*\*** | **,801\*\*** | ,415 | **,535\*** | ,400 | ,320 | ,156 | ,317 | -,118 |
|  | Sig. | **,004** | **,001** | **,001** | **,019** | **,034** |  | **,000** | **,000** | **,000** | **,000** | ,069 | **,015** | ,081 | ,168 | ,510 | ,173 | ,620 |
| *PASS-C* | Corr.coeff. | **,503\*** | **,559\*** | **,559\*** | **,462\*** | ,268 | **,895\*\*** | 1,000 | **,737\*\*** | **,690\*\*** | **,707\*\*** | ,329 | ,424 | ,344 | ,227 | ,118 | ,214 | -,254 |
|  | Sig. | **,024** | **,010** | **,010** | **,040** | ,254 | **,000** |  | **,000** | **,001** | **,000** | ,157 | ,063 | ,137 | ,336 | ,621 | ,366 | ,280 |
| *PASS-E* | Corr.coeff. | **,614\*\*** | **,777\*\*** | **,766\*\*** | **,529\*** | **,564\*\*** | **,899\*\*** | **,737\*\*** | 1,000 | **,918\*\*** | **,544\*** | **,472\*** | **,592\*\*** | ,419 | ,387 | ,161 | ,330 | -,062 |
|  | Sig. | **,004** | **,000** | **,000** | **,016** | **,010** | **,000** | **,000** |  | **,000** | **,013** | **,036** | **,006** | ,066 | ,092 | ,499 | ,156 | ,795 |
| *PASS-F* | Corr.coeff. | **,556\*** | **,666\*\*** | **,643\*\*** | **,502\*** | **,486\*** | **,886\*\*** | **,690\*\*** | **,918\*\*** | 1,000 | **,541\*** | **,577\*\*** | **,736\*\*** | **,486\*** | ,291 | ,188 | **,445\*** | ,085 |
|  | Sig. | **,011** | **,001** | **,002** | **,024** | **,030** | **,000** | **,001** | **,000** |  | **,014** | **,008** | **,000** | **,030** | ,213 | ,428 | **,049** | ,720 |
| *PASS-P* | Corr.coeff. | **,647\*\*** | **,558\*** | **,565\*\*** | **,502\*** | ,421 | **,801\*\*** | **,707\*\*** | **,544\*** | **,541\*** | 1,000 | ,261 | ,304 | ,328 | ,289 | ,112 | ,118 | -,023 |
|  | Sig. | **,002** | **,011** | **,009** | **,024** | ,065 | **,000** | **,000** | **,013** | **,014** |  | ,267 | ,193 | ,157 | ,216 | ,639 | ,619 | ,925 |
| *FABQ* | Corr.coeff. | ,337 | ,344 | ,360 | ,351 | ,336 | ,415 | ,329 | **,472\*** | **,577\*\*** | ,261 | 1,000 | **,781\*\*** | **,951\*\*** | ,314 | -,032 | ,195 | ,009 |
|  | Sig. | ,146 | ,138 | ,120 | ,130 | ,147 | ,069 | ,157 | **,036** | **,008** | ,267 |  | **,000** | **,000** | ,178 | ,894 | ,410 | ,970 |
| *FABQ-PA* | Corr.coeff. | **,494\*** | **,442\*** | ,404 | ,315 | ,375 | **,535\*** | ,424 | **,592\*\*** | **,736\*\*** | ,304 | **,781\*\*** | 1,000 | **,638\*\*** | ,140 | ,009 | ,377 | ,178 |
|  | Sig. | **,027** | **,050** | ,077 | ,176 | ,103 | **,015** | ,063 | **,006** | **,000** | ,193 | **,000** |  | **,002** | ,557 | ,969 | ,101 | ,452 |
| *FABQ-W* | Corr.coeff. | ,280 | ,339 | ,378 | ,411 | ,280 | ,400 | ,344 | ,419 | **,486\*** | ,328 | **,951\*\*** | **,638\*\*** | 1,000 | ,291 | -,069 | ,185 | -,040 |
|  | Sig. | ,231 | ,144 | ,101 | ,071 | ,231 | ,081 | ,137 | ,066 | **,030** | ,157 | **,000** | **,002** |  | ,213 | ,772 | ,435 | ,867 |
| *S-ANXIETY* | Corr.coeff. | **,449\*** | **,451\*** | **,470\*** | **,629\*\*** | ,284 | ,320 | ,227 | ,387 | ,291 | ,289 | ,314 | ,140 | ,291 | 1,000 | ,128 | -,198 | ,090 |
|  | Sig. | **,047** | **,046** | **,037** | **,003** | ,224 | ,168 | ,336 | ,092 | ,213 | ,216 | ,178 | ,557 | ,213 |  | ,592 | ,402 | ,707 |
| *T-ANXIETY* | Corr.coeff. | ,299 | ,139 | ,071 | ,034 | ,035 | ,156 | ,118 | ,161 | ,188 | ,112 | -,032 | ,009 | -,069 | ,128 | 1,000 | ,185 | ,378 |
|  | Sig. | ,200 | ,558 | ,766 | ,886 | ,883 | ,510 | ,621 | ,499 | ,428 | ,639 | ,894 | ,969 | ,772 | ,592 |  | ,435 | ,100 |
| *Rating harmful* | Corr.coeff. | ,133 | ,240 | ,276 | ,044 | ,270 | ,317 | ,214 | ,330 | **,445\*** | ,118 | ,195 | ,377 | ,185 | -,198 | ,185 | 1,000 | ,289 |
|  | Sig. | ,577 | ,307 | ,238 | ,854 | ,250 | ,173 | ,366 | ,156 | **,049** | ,619 | ,410 | ,101 | ,435 | ,402 | ,435 |  | ,217 |
| *Rating harmless* | Corr.coeff. | ,289 | ,168 | ,091 | ,178 | ,135 | -,118 | -,254 | -,062 | ,085 | -,023 | ,009 | ,178 | -,040 | ,090 | ,378 | ,289 | 1,000 |
|  | Sig. | ,216 | ,479 | ,703 | ,452 | ,570 | ,620 | ,280 | ,795 | ,720 | ,925 | ,970 | ,452 | ,867 | ,707 | ,100 | ,217 |  |

**Table 3.**
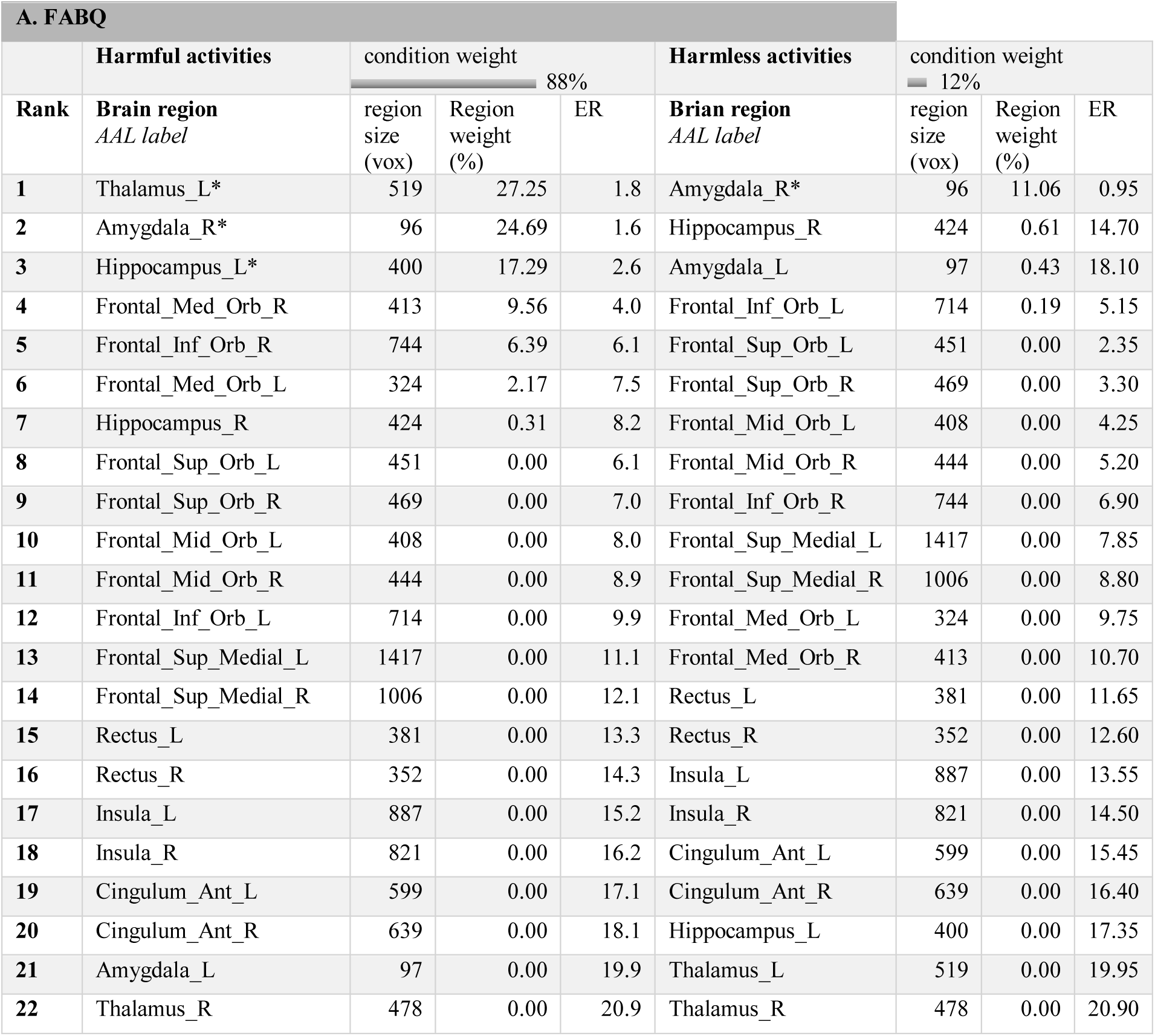
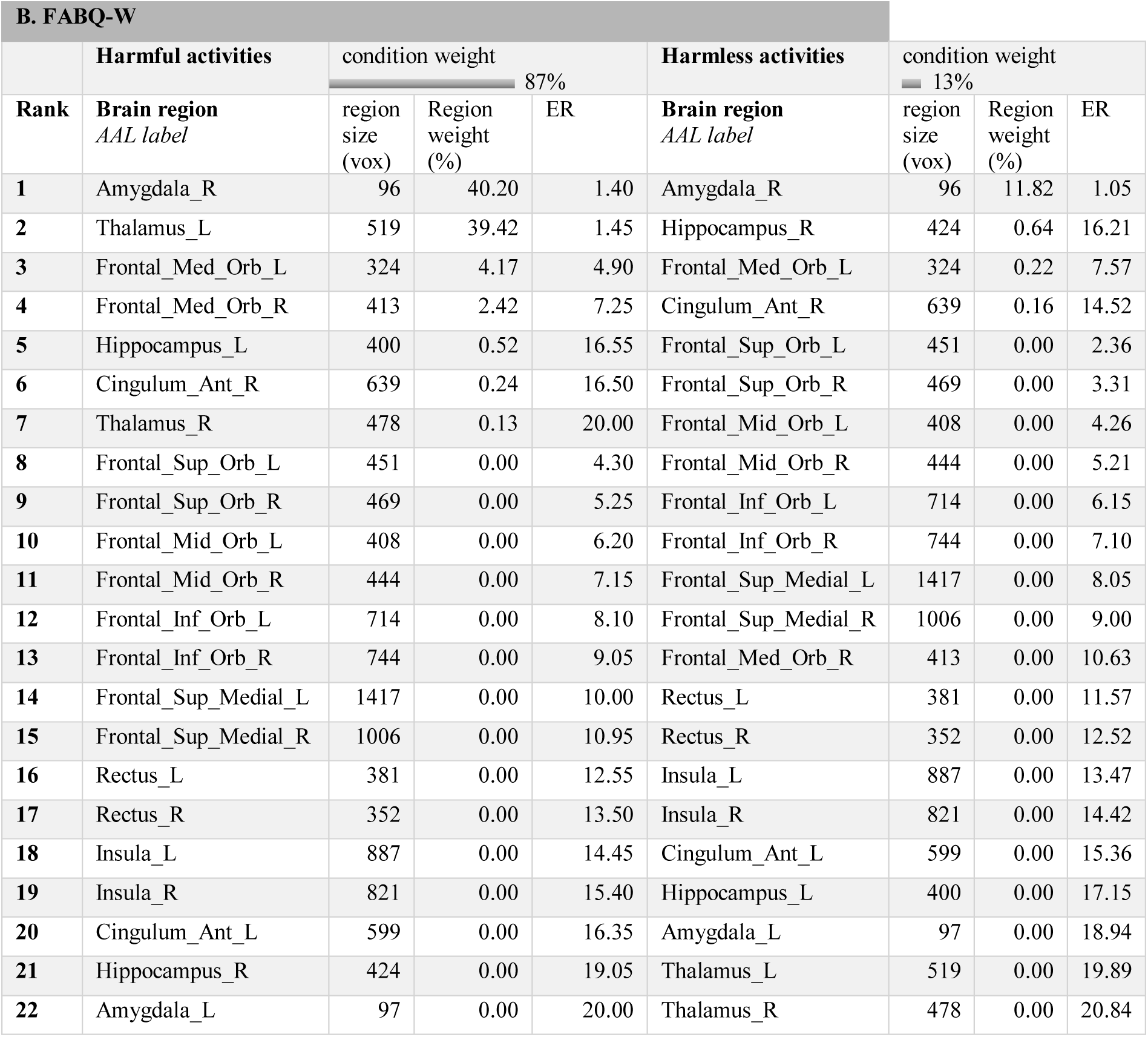
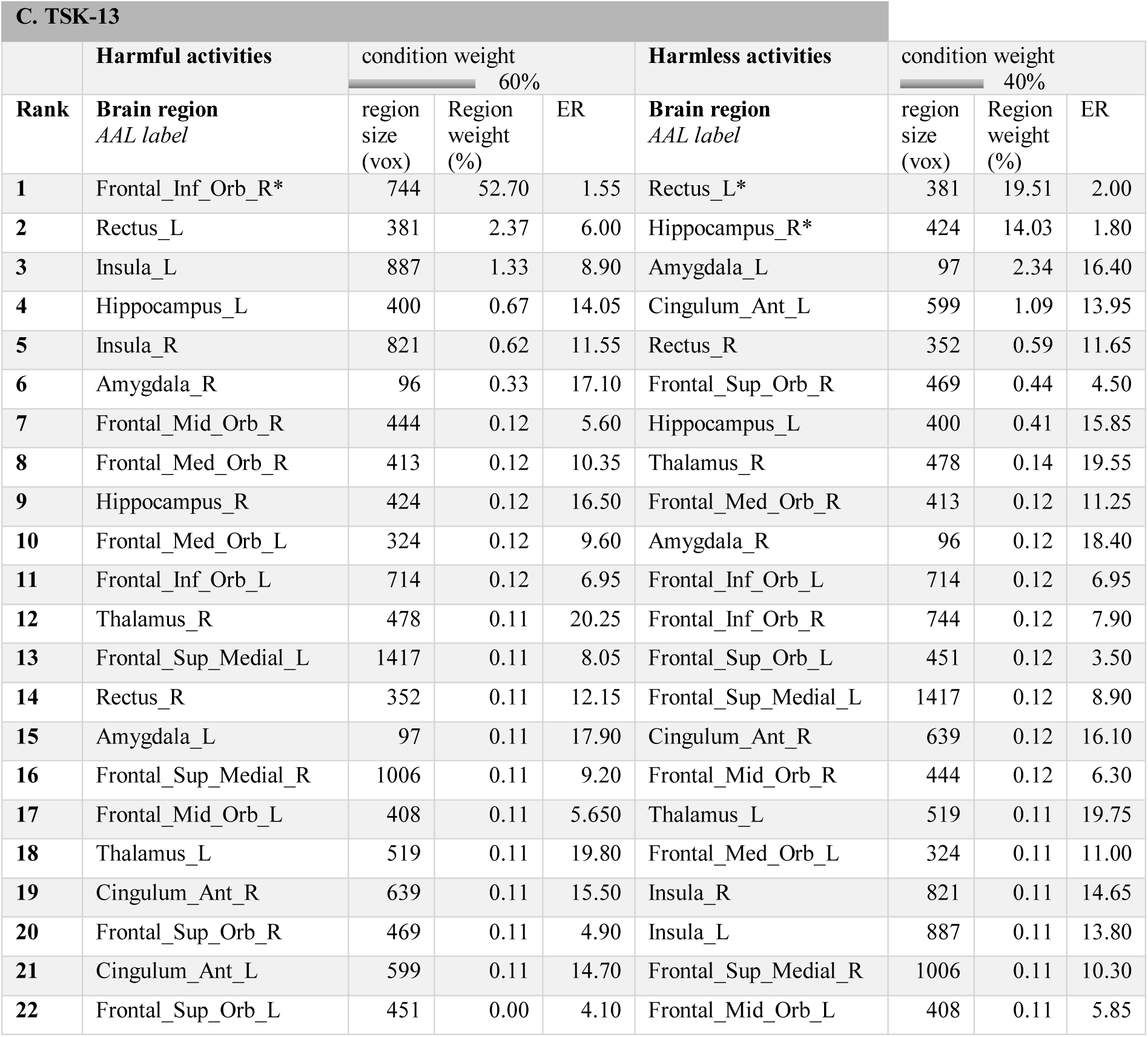
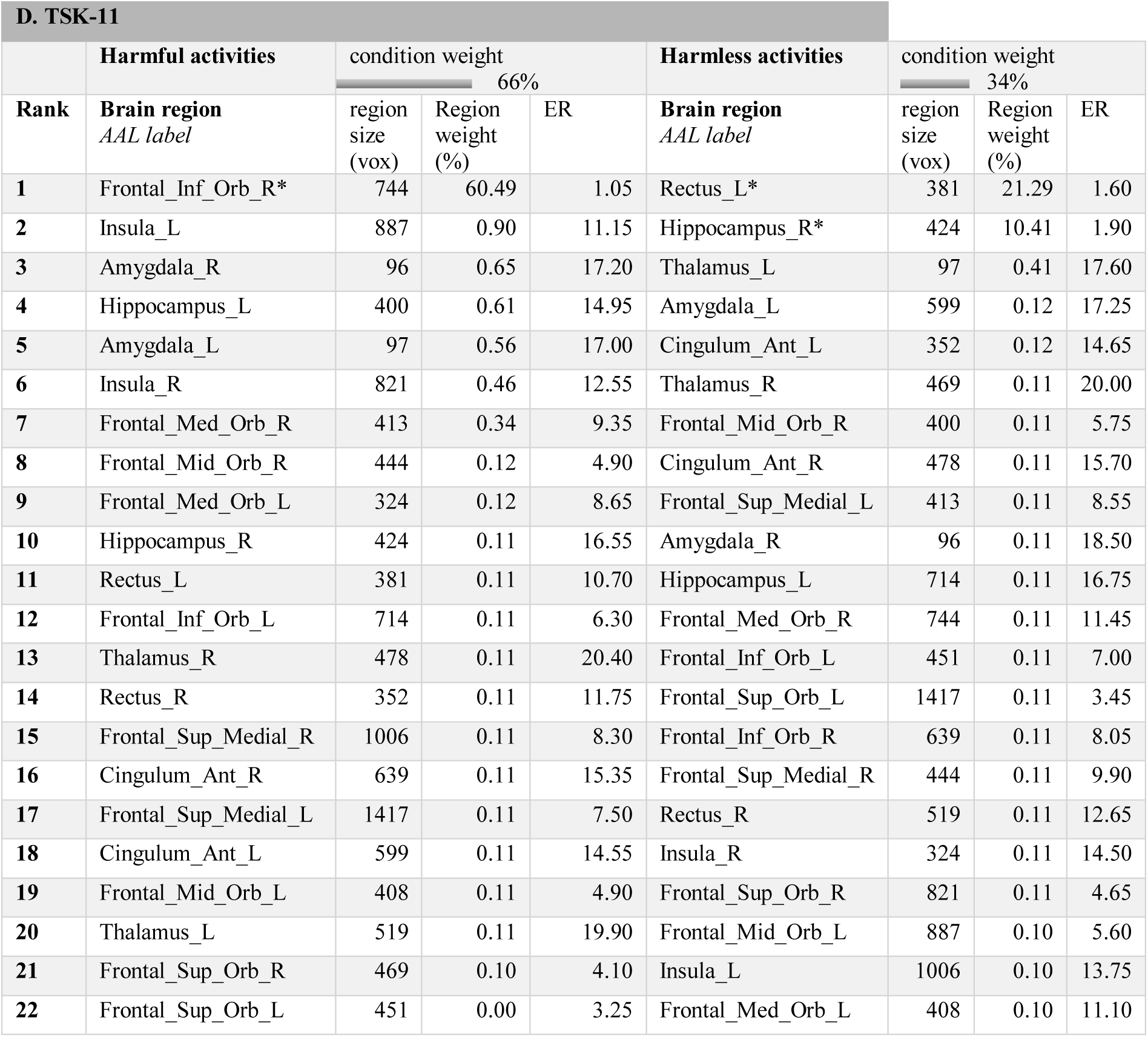
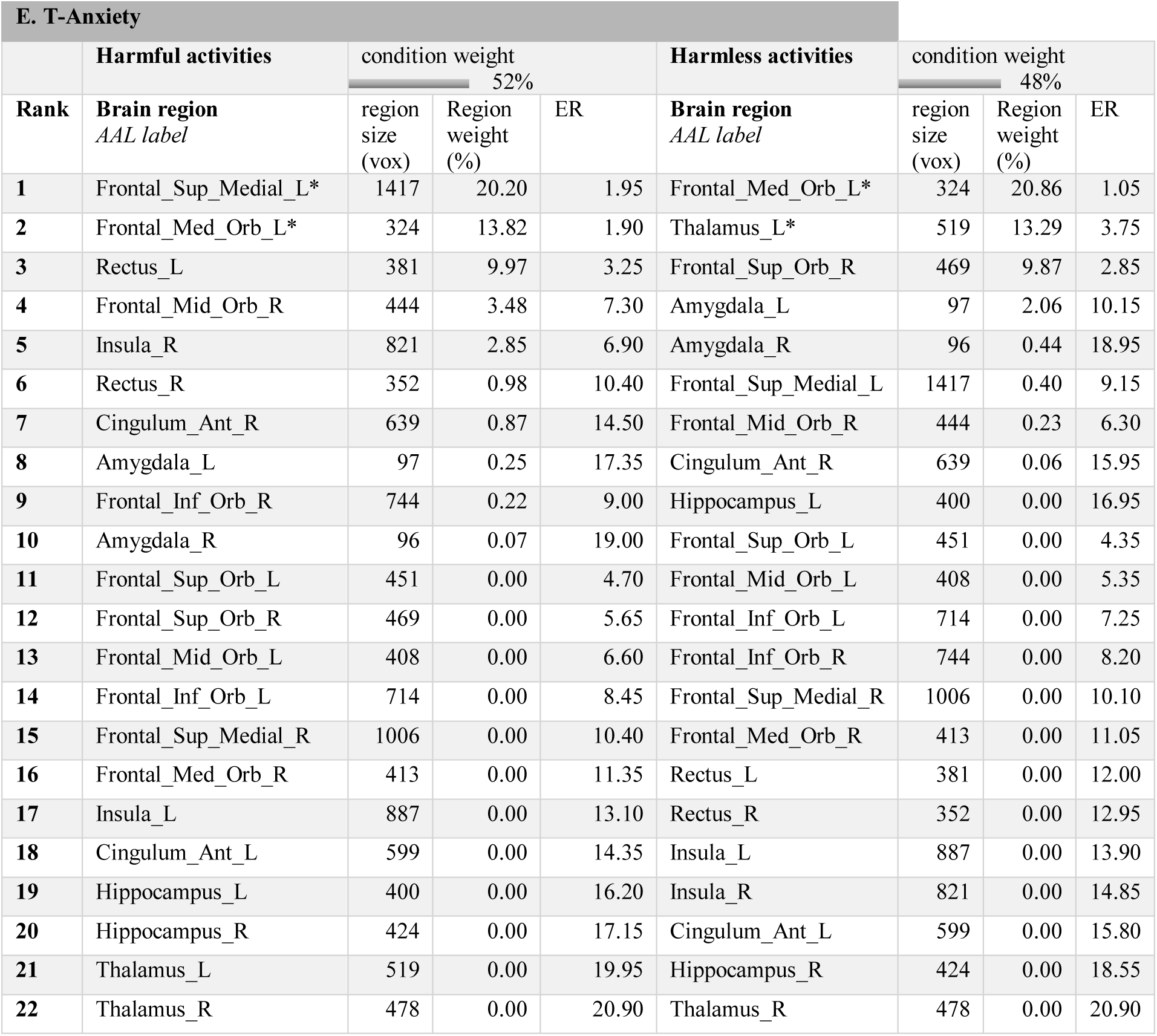
Condition and region weights showing the contribution of the two different conditions and fear-related brain regions to the final decision function of each MKL model (questionnaires A-E with model performance p < 0.05) in hierarchical order. The brain regions (left and right hemisphere) were parcellated according to the AAL atlas: Medial orbitofrontal regions (mOFC: Rectus, Frontal_Sup_Orb, Frontal_Med_Orb), lateral orbitofrontal regions (lOFC: Frontal_Mid_Orb, Frontal_Inf_Orb), medial prefrontal cortex (mPFC: Frontal_Sup_Medial), anterior cingulate cortex (Cingulum_Ant), Thalamus, Amygdala, Hippocampus and Insula. ER = expected ranking. *predictive brain set for secondary cross-validation

### 2.7 Definition of brain regions and atlas registration

Based on a-priori knowledge of brain regions involved in fear processing, we limited the feature space to bilateral fear-related brain regions including the amygdala, hippocampus, thalamus, anterior cingulate, insula, medial prefrontal and orbitofrontal cortices (Tovote et al., 2015; Braem et al., 2017; Meier et al., 2014). The respective brain regions were parcellated according to the Automated Anatomical Labeling (AAL, see Table 3 for the different labels) (Tzourio-Mazoyer et al., 2002) atlas and projected on the ICBM 152 Nonlinear template (section 2.4) by means of MATLAB (version R2017b) based surface-volume registration tools (svreg) implemented in BrainSuite (version 17a) (Shattuck and Leahy, 2002). BrainSuite was also used to generate surfaces of the selected AAL regions for visualization.

### 2.8 Model evaluation and interpretation

Model performance was assessed by two metrics commonly used to assess the performance of regression models (Ivanescu et al., 2016; Fernandes et al., 2017): Pearsons’s correlation coefficient (r) and the mean squared error (MSE). The correlation coefficient characterizes the linear relationship between observed and predicted labels; the MSE is calculated as the average of the squared differences between the observed and predicted labels. A significant positive correlation between observed and predicted labels would indicate strong decoding performance. Unlike in conventional correlation analysis, however, a negative correlation would indicate poor performance. For each model, we report the normalized MSE (nMSE) because the different questionnaires are based on different score ranges. To explore possible differential contributions of fear-related brain regions to the prediction models, we report the contribution rank of each brain region (region weight) within each condition (condition weight) provided by the MKL approach (Table 3). Importantly, the selection of regions by the MKL model might be influenced by small variations in the dataset (as induced by cross-validation) and might therefore lead to different subsets of regions being selected across cross-validation steps (folds). Providing a quantification of this variability, the “expected ranking (ER)” (see Table 3) characterizes the stability of the region ranking across folds: The closer the ER to the ranking of the selected fold, the more consistent is the ranking of the respective brain region across folds. On the other hand, if the ER is different from the ranking, this means that the ranking might be variable across folds.

## 3. Results

### 3.1 Ratings, questionnaire scores and correlations

Importantly, the comparison of the ratings during fMRI measurements demonstrated that the potentially harmful activities were perceived as being significantly more harmful compared to the harmless activities (paired-T-Test: T = 8.22, p < 0.001, two-tailed). Descriptive statistics of the different questionnaires as well as age and sex of the patients are summarized in Table 1. Regarding the questionnaire data, visual inspection and the Shapiro-Wilk test indicated non-normality of the data (p<0.05) of several questionnaires (FABQ, FABQ-W, FABQ-PA and T-Anxiety) and therefore, the non-parametric Spearman’s rank correlation coefficient was used.

Several significant positive correlations between the different PRF questionnaires scores were observed (p < 0.05, Table 2). Most of the TSK scales significantly correlated with the PASS scales (0.97 < r’s > 0.46, p < 0.05) whereas the FABQ work scale did not show significant relationships with the TSK and PASS scales (p > 0.05), except for the PASS-F scale (r = 0.49, p < 0.05). Furthermore, only the S-Anxiety scale of the STAI scale demonstrated significant correlations with some, but not all, TSK scales (0.44 < r’s > 0.63, p < 0.05). Finally, only the PASS-F scale showed a positive and significant relationship with the mean rating of the harmful condition (r = 0.44, p < 0.05, Table 2).

### 3.2 Model performance

The MKL models with significant performance results (p < 0.05) characterized by the Pearsons’s correlation coefficient (r) and the normalized mean squared error (nMSE) are depicted in Figure 1 (A-E). The FABQ model demonstrated a significant decoding performance characterized by a positive correlation between true and predicted labels (r = 0.61, nMSE =4.25, p < 0.05). Interestingly, the FABQ-W model showed strong predictive power (r = 0.74, nMSE = 1.81, p < 0.05) whereas the FABQ-PA scale was not decodable from fear-related brain response patterns (r = 0.03, nMSE = 1.68, p > 0.05). Among the TSK scales, only the TSK-13- (r = 0.37, nMSE = 1.09, p < 0.05) and the TSK-11- (r = 0.60, nMSE = 0.90, p < 0.05) models demonstrated a significant decoding performance. The TSK-17 model (r = 0.19, nMSE = 1.10, p > 0.05) and the TSK-11 subscale models did not show a significant decoding performance (TSK11-SF: (r = −0.73, nMSE = 0.86, p > 0.05 / TSK-11-AA: r = −0.63, nMSE = 0.88, p > 0.05). In addition, none of the PASS scales were decodable from fear-related brain response patterns (p’s > 0.05, PASS: r = 0.18, nMSE = 4.63 / PASS-C: r = −0.44, nMSE = 1.64 / PASS-E: r = −0.32, nMSE = 1.38, PASS-F: r = −0.15, nMSE = 1.70 / PASS-P: r = −0.51, nMSE = 1.36). Furthermore, and interestingly, the T-Anxiety model demonstrated a moderate decoding performance (r = 0.48, nMSE = 1.01) whereas the S-Anxiety model was not significant (r = −0.46, nMSE = 1.51, p > 0.05). Finally, the ratings of perceived harmfulness during fMRI measurements were not decodable from fear-related brain response patterns (Rating harmful: r = −0.01, nMSE = 0.64, p > 0.05 / Rating harmless: r = −0.72, nMSE = 0.38, p > 0.05).

**Figure 1.**
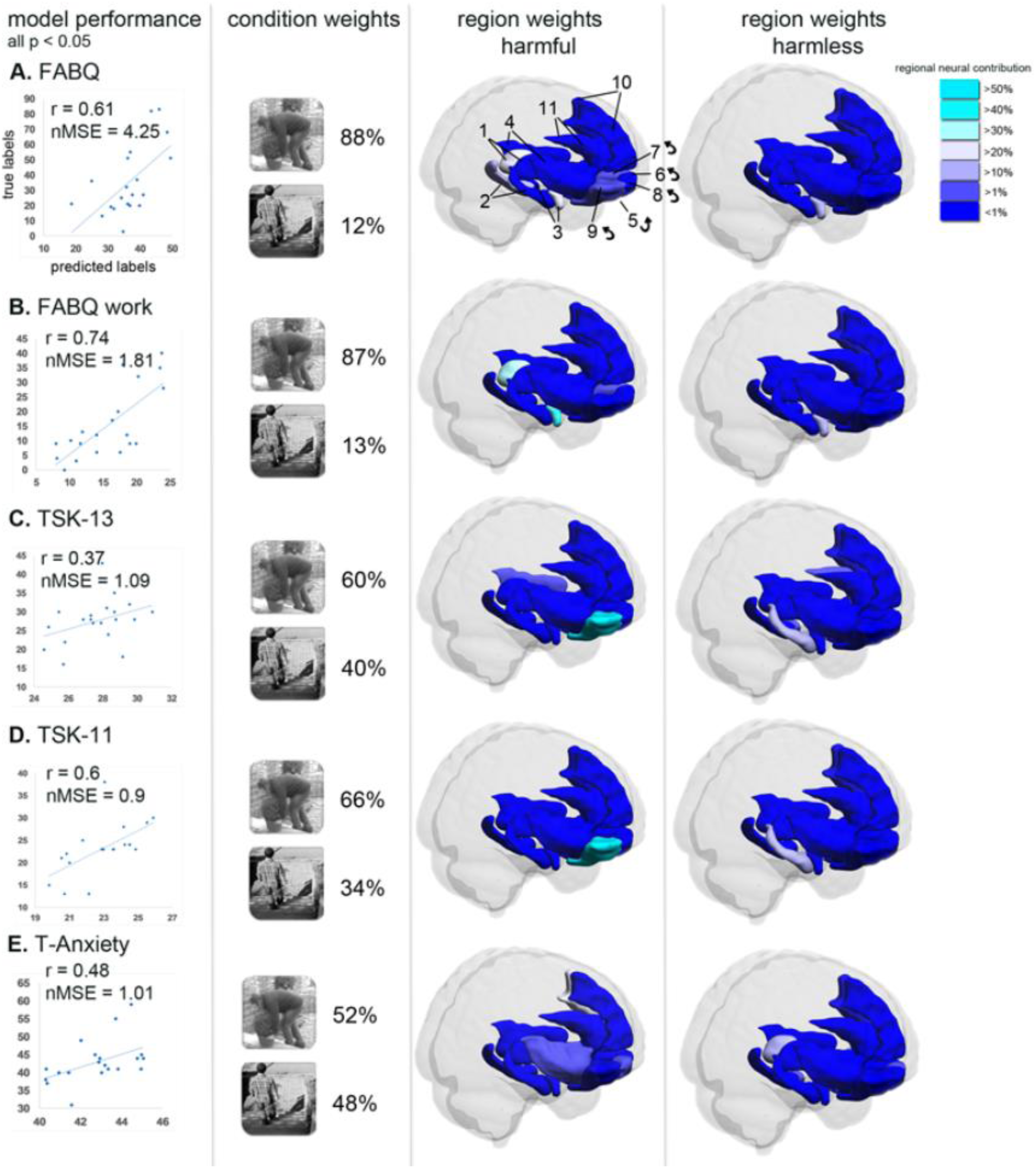
The model performance (r, MSE) characterizes the strength of relationship between true and predicted labels. Condition and region weights show the predictive contribution of the two different conditions (harmful, harmless) and fear-related brain regions (parcellated according to the AAL atlas, L = left, R = right) to the final decision function of each MKL model (questionnaires A-E with model performance p < 0.05). Brain regions (feature set): Thalamus (1), Hippocampus (2), Amygdala (3), Insula (4), mOFC: Rectus (5), Frontal_Sup_Orb (6), Frontal_Med_Orb (7), lateral OFC: Frontal_Mid_Orb (8), Frontal_Inf_Orb (9)), mPFC: Frontal_Sup_Medial (10), anterior cingulate cortex (Cingulum_Ant (11)). 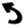 indicates not visible contralateral homologue.

### 3.3 Condition and region weights

The condition and region weights of models with significant performance (p<0.05, section 3.2) are illustrated in Figure 1 (A-E) and described in detail in Table 3 (A-E). The decoding performances of the FABQ models (FABQ and FABQ-W) were driven by a major contribution of the harmful condition (88% and 87%, respectively). Within this condition, the left thalamus (rank 1), the right amygdala (rank 2) and the left hippocampus (rank 3) contributed more than 69% of the total region weights in the FABQ model (Table 3A). Similarly, the right amygdala (rank 1) and the left thalamus (rank 2) carried the most predictive neural information with 79,42% of the total region weights in the FABQ-W model (Table 3B). In both FABQ models, the right amygdala also demonstrated an association with the harmless condition, although of minor relevance (∼11%). By comparison, the TSK models demonstrated a moderate contribution of the harmful condition (TSK-13:60%, TSK-11:66%). Both predictive model performances of the TSK were driven by a major contribution of the right lateral orbitofrontal cortex (lOFC, TSK-13: 52.7%, TSK-11: 60,49%, Table 3C, 3D). Furthermore, the left medial orbitofrontal cortex (mOFC) and the right hippocampus carried predictive information within the harmless condition in both TSK models (TSK-13: left gyrus rectus 19.51%, right hippocampus: 14.03% / TSK-11: left gyrus rectus: 21.29%, right hippocampus: 10.41%). Interestingly and with almost equal contributions of the harmful (52%) and harmless conditions (48%), the predictive model of T-Anxiety was mainly driven by neural contributions of the left medial prefrontal (mPFC) and mOFC (accounting for 44% of the total region weights in the harmful condition) and the left thalamus (together with the mOFC accounting for 44% of the total region weights in the harmless condition, Table 3E). Finally, the secondary cross-validation using each predictive brain set of the significant models (FABQ, TSK-13, TSK-11, T-Anxiety) and training and testing it with the labels of the other questionnaires did not result in significant performance results (p’s > 0.05).

## 4. Discussion

Evidence from cross-sectional and longitudinal behavioral studies demonstrates a strong association between PRF and disability in chronic pain (Leeuw et al., 2007b; Esteve et al., 2017; Wertli et al., 2014b). However, the different PRF constructs such as “fear of movement/(re)injury/kinesiophobia”, “fear avoidance beliefs” or “pain anxiety” are often used interchangeably in the literature (Lundberg et al., 2011) and it is unclear if they share a common PRF construct reflected by similar neural sources. The (sub-)cortical neural basis of fear and anxiety that controls cognitions and regulates appropriate behavior dependent on threat characteristics is well described (Gray and MacNaughton, 2000; McNaughton and Corr, 2004; Qi et al., 2018; Shackman et al., 2011; Panksepp, 2011; LeDoux, 2000). Altough both emotions are linked to similar neuromodulatory systems of the fear circuit (Tovote et al., 2015), anxiety is less well understood and more complex than fear. Current research suggests a functional differentiation characterized by subcortical regions processing fast fear responses to an imminent threat (defensive responses) and cortical systems processing complex cognitions related to fear and anxiety where the threat is distal in space or time (Qi et al., 2018; LeDoux and Pine, 2016).

The current MVPA approach using MKL demonstrated the feasibility to neuronally dissect the proposed constructs of PRF self-reports based on their (sub-)cortical predictors during PRF related brain activity. The results revealed that while the individual variability of some questionnaires, specifically the FABQ- and FABQ-W-, TSK-13, TSK-11 and T-Anxiety-scale, was predictable from response patterns in fear-related, dissociable neural sources on subcortical and cortical levels, this was not the case for the FABQ-PA-, the TSK-11 subscales (TSK-11-AA and TSK-SF), the PASS scales and the S-Anxiety scale. Furthermore, the online ratings of perceived harmfulness were not decodable from fear-related brain response patterns.

### FABQ and TSK

The FABQ and FABQ-W scales demonstrated the best model performances among the investigated PRF questionnaires, characterized by a strong contribution of neural information in the harmful condition (condition weights: 88% and 87%, respectively). Interestingly, the FABQ-PA scale did not show a predictive association with fear-related brain response patterns. The better model performance of the FABQ-W is in line with the emerging evidence that the FABQ-W is a stronger moderator of treatment efficacy in chronic LBP compared to the FABQ-PA, although this might be dependent on the patient population (George et al., 2005; George et al., 2008; Waddell et al., 1993; Wertli et al., 2014a). In support of this, the FABQ-W scale qualified for a clinical prediction rule regarding improvement after spinal manipulation, whereas the FABQ-PA scale did not (Dougherty et al., 2014; Flynn et al., 2002).

With respect to the region weights, the FABQ models were mainly driven by subcortical neural contributions involving the thalamus, hippocampus and the amygdala while frontal brain regions played a minor role. The thalamus and particularly its midline structures have been considered to be a non-specific arousing system (van der Werf et al., 2002). However, it has been recently shown that parts of dorsal midline thalamic structures are necessary for fear memory processing by directly targeting the hippocampus, which plays an important role for context-dependent emotional memory (Penzo et al., 2015; Lara-Vásquez et al., 2016; Zheng et al., 2017). On the other hand, the amygdala has long been considered a “fear center” (Panksepp, 1998; Darwin, 1873). However, the heterogeneous structure consisting of several nuclei is not essential for the *experience* of fear, demonstrated in patients with amygdala lesions (Anderson and Phelps, 2002; Feinstein et al., 2013; LeDoux and Pine, 2016). Instead, the amygdala has been shown to be more strongly implicated in behavioral and physiological responses to threats (i.e. defensive processes); its relation to complex cognitions like fear and anxiety is controversial (Fanselow and Pennington, 2017; LeDoux and Pine, 2016; Panksepp, 2011). A recent opinion paper suggests that subjective feelings of fear and anxiety do not initially arise from subcortical activity of the fear circuit centered around the amygdala (LeDoux and Hofmann, 2018). Thus, amygdala activity and mediated physiological responses of fear and anxiety might be, at its best, only a correlate of subjective feelings of fear and anxiety (LeDoux and Hofmann, 2018). However, and in contrast to this view, the results presented here indicate a strong predictive association between subjective reports of PRF, assessed by the FABQ scales, and the amygdala.

Among the TSK scales, the TSK-13 and the TSK-11 demonstrated a predictive association with fear-related brain response patterns, albeit with less contribution of the harmful condition compared to the FABQ scales (TSK-13: 60% and TSK-11: 66%). The TSK-11 version showed a stronger relationship between true and predicted labels compared to the TSK-13 version (r = 0.60, nMSE = 0.90, p < 0.05). This result might reflect the progress of previous research regarding the psychometric properties of the different TSK versions. Compared to the 17-item version, the 13-item version has better psychometric properties without the four inversely phrased items (Roelofs et al., 2004a; Neblett et al., 2016) and the 11-item version has been recommended for future research and clinical settings (for a chronological summary see Tkachuk and Harris, 2012). Interestingly, no predictive association could be “learned” by MKL using the TSK-11 subscale labels (TSK-11-SF and TSK-11-AA scores). Although these two lower order factors (activity avoidance and somatic focus) are reflective of the higher order construct “fear of movement and (re)injury/kinesiophobia”, the non-significant result might indicate that they are associated with inconsistent neural patterns across individuals.

Regarding the region weights of the TSK models, the right lateral orbitofrontal cortex (lOFC) provided the most predictive information for the two TSK scales (TSK-13: 52%, TSK-11: 60%). In agreement with the phobia-related construct (kinesiophobia), dysfunction of the OFC has been shown to be implicated in the processing of phobia-related stimuli in disorders such as social anxiety disorder (Dilger et al., 2003). Specifically, lOFC activity was reduced when phobogenic trials were contrasted with fear-relevant trials (Aue et al., 2015). Furthermore, a hyperactive lOFC has been shown to be linked to anxiety-laden cognitions (Hahn et al., 2011). Interestingly, the higher cortical contributions of the TSK models were clearly dissociable from the largely subcortical contributions involving the amygdala, hippocampus and thalamus that predicted the FABQ scores.

To conclude, the FABQ scales demonstrated high PRF sensitivity (harmful condition weights > 87%) and were linked to subcortical predictors that have been associated with fear responses to an imminent threat and defensive behavior (LeDoux and Pine, 2016; McNaughton and Corr, 2004). In contrast, the TSK scales appeared to capture emotional states largely associated with cortical fear processing that might be related to cognitive aspects of PRF. In support of this, the observed higher harm*less* condition weights of the TSK compared to the FABQ models might indicate that the TSK scales are associated with more diffuse anxiety-related cognitions.

### PASS

Surprisingly, the PASS failed to demonstrate a predictive association with fear-related brain response patterns. There may be several explanations. First, whereas the FABQ and the TSK scales have been specifically developed for patients with musculoskeletal pain, the PASS is suitable for various pain phenotypes (Crombez et al., 1999). Second, the PASS has been shown to be more strongly associated with negative affect and was less predictive of pain disability and behavioral performance (Crombez et al., 1999). Third, in a recent study assessing fear of bending, the PASS (and the TSK) score was not related to physiological measurements such as startle responses (Caneiro et al., 2017). Fourth, all PASS subscales demonstrated significant multicollinearity in our sample suggesting non-independence between the different subscales. All these aspects may have led to less sensitivity of fear related neural patterns to the PASS and its subscales in the current study.

The superiority of the FABQ scale (driven by the FABQ-W) in decoding performance compared to the TSK and PASS scales might also be influenced by the back-specific items of the FABQ in conjunction with the nature of the PRF-provoking stimuli (back straining movements). The items of the FABQ were specifically related to the back while the TSK and PASS can be used with various musculoskeletal pain diagnoses such as work-related upper extremity disorders, chronic LBP, fibromyalgia, and osteoarthritis (Roelofs et al., 2007). Nevertheless, the FABQ has also been adapted to shoulder pain where it demonstrated better factor structure and a stronger association with disability compared to the TSK-11 (Mintken et al., 2010).

### State and Trait anxiety

Beside PRF, anxiety and depression significantly mediate the relationship between pain and disability (Marshall et al., 2017). Nevertheless, fear responses specifically related to a patient’s pain and/or potentially painful movements might be more relevant for explaining disability in chronic LBP than general trait anxiety responses (McCracken et al., 1996). The current results are in line with this notion. First, most of the PRF measures did not show a significant relationship with state or trait anxiety. Second, state anxiety was not decodable from fear-related brain responses to potentially harmful activities in chronic pain patients. Interestingly, with respect to the trait anxiety model (T-Anxiety, Figure 1E), the harmful (52%) and the harmless conditions (48%) carried almost equal predictive neural information. This suggests that the trait anxiety measure is associated with neural content irrespective of the harmfulness of a stimulus, provoked by e.g. enhanced attention to visual information processed in fear-related brain regions (Berggren et al., 2015). This might indicate that the T-Anxiety scale captures neural responses that are associated with a more generalized fear response. This notion is supported by a study showing that individuals with high trait anxiety exhibit sustained PRF during extinction (Meulders et al., 2014).

Regarding the regions weights, predictive information was predominantly provided by brain regions that were less involved in the prediction of the other PRF measures, namely parts of the mPFC and mOFC (Table 3E). This is in line with the proposed functional differentiation of neural structures regarding fear in response to an imminent threat (defensive response) and cognitive fear/anxiety (distal, uncertain threat) whereas the latter involves more rostral cortical structures such as the mPFC and mOFC (LeDoux and Pine, 2016; McNaughton and Corr, 2004). Moreover, research on self-report measurements indicates that trait anxiety is relatively distinct from tissue damage fear, which supports a behavioral and neural dissociation of trait anxiety and PRF (Perkins et al., 2007; Cooper et al., 2007).

### Harmfulness ratings

Interestingly, although the PRF-provoking harmful activities were significantly rated as more harmful compared to the harmless activities, the ratings of perceived harmfulness during fMRI measurements were not decodable from fear-related brain response patterns. Furthermore, the ratings did not show significant correlations with PRF measures (except the PASS-F scale, see Table 2). Others reported only moderate relationships (r’s < 0.39) between perceived harmfulness ratings of PHODA items and self-report measures such as the TSK, Pain Catastrophizing scale (PCS) or pain intensity (Leeuw et al., 2007a), indicating that ratings of perceived harmfulness assess something akin to, but also distinct from PRF self-report measures. The weak relationships between ratings of perceived harmfulness and self-report measures of PRF might be explained by the specificity of the potentially harmful movements depicted by the PHODA items. Namely, the ratings of perceived harmfulness were specifically related to back straining movements such as bending and lifting while the PRF measures might also be associated with other potentially harmful movements. As such, the fear-related neural patterns induced by the observation of potentially harmful activities for the back might not include information about movement specificity. Instead, these neural patterns might predict PRF and its constructs in a more general fashion that is captured by the TSK and FABQ.

### Limitations

A limitation of this study is the relatively small sample size in conjunction with the cross-validation framework. Ideally, the predictive model should be trained and tested with completely independent data. However, the results obtained are likely to be valid for several reasons: 1) the goal of the current study was not maximizing decoding performance, rather, multivariate decoding was used for the interpretation and understanding of the different fear constructs, for which significant predictive accuracy was obtained (Hebart and Baker, 2017); 2) the applied linear support vector machines have been shown to exhibit good performance even in very high dimensional settings with small sample sizes (Varoquaux and Thirion, 2014); 3) the applied regression approach using continuous variables enhances statistical power compared to a categorical analysis (e.g. low versus high fear) (Altman and Royston, 2006); 4) the variability of the regions most contributing to the models across cross-validation folds was very small (indicated by the expected ranking (ER)), demonstrating stable ranking across folds. For these reasons, the differences of the prediction models are unlikely to be caused by the small sample size. Finally, the study design only allows interpretations of PRF to back straining movements and LBP. Therefore, conclusions related to other musculoskeletal conditions should be drawn with caution. Nevertheless, the current approach might represent a promising new tool to dissect psychological constructs of self-report measures.

### Conclusion

This is the first time that multivariate brain responses patterns are used to better understand and dissect a psychological construct, here, PRF, conventionally assessed by self-report (questionnaires). The FABQ scale demonstrated strong predictive power with high sensitivity to the harmful condition and was associated with subcortical fear processing regions (amygdala, thalamus, hippocampus). The TSK scales were more related to neural content of higher order structures such as the OFC and showed less sensitivity to the harmful condition compared to the FABQ scales. This might indicate that the construct of kinesiophobia is more related to diffuse anxiety related neural systems whereas the FABQ scales are more related to defensive systems of fear. Finally, the PASS and its subscales failed to demonstrate a predictive association with fear-related brain response patterns. To conclude, while self-reports still represent the best and direct measure of subjective feelings of fear and anxiety (LeDoux and Hofmann, 2018), the current results emphasize the need to carefully consider the different PRF questionnaires in research and clinical settings as their constructs are not interchangeable.

## Acknowledgements

We would like to thank Dr. Nina Kreddig (Ruhr University Bochum) and Dr. Stefan Sommer (ETH Zurich) for their intellectual contributions. Finally, we thank Sergio Maffioletti from S3IT (University of Zurich) for technical support regarding the supercomputing environment.

